# Fitness flux in SARS-CoV-2 and influenza H3N2

**DOI:** 10.64898/2026.07.05.736619

**Authors:** Trevor Bedford

## Abstract

The tempo of viral adaptation is usually read indirectly from the composition of mutations, through measures such as dN/dS. Here we measure it directly from the dynamics of variant frequencies, where we use multinomial logistic regression to estimate a fitness for each co-circulating variant. We aggregate these estimates to derive the rate of change of mean population fitness, referred to as fitness flux. Tracing SARS-CoV-2 from its emergence, we find that it initially adapted rapidly, doubling in fitness every 6 months from Jan 2021 to Jun 2022, but slowing to every 2.4 years from Jul 2022 to Dec 2025. Seasonal influenza H3N2 sustained a slower, steadier pace doubling in fitness every 10.0 years. In both, the rate of fitness gain closely tracks the variance in fitness, matching the 1:1 expectation of Fisher’s fundamental theorem. Phylogenetic contrasts between parent and child lineages localize most fitness gain to spike, and within spike to the receptor-binding domain, where a simple count of spike S1 substitutions predicts lineage fitness about as well as deep-learning escape and protein-language-model scores. Measuring fitness directly thus offers a transparent, frequency-based alternative to mutational proxies for tracking and anticipating viral adaptation.

*The website blab.github.io/fitness-flux/ is the intended reading experience of this paper, providing responsive layout and interactive figures*.

## Introduction

RNA viruses evolve rapidly and the selective pressures they face shift over the course of emergence to endemicity. A newly established virus initially adapts to its new host by refining its capacity for within-host replication and between-host transmission. Once a virus becomes endemic, adaptation is instead dominated by continual escape from accumulating population immunity, driving ongoing antigenic change, where some viruses are better able than others to sustain adaptive evolution [1]. Adaptation of either kind leaves a signature in amino-acid replacing nonsynonymous vs silent synonymous substitutions, with methods ranging from simple comparisons of nonsynonymous to synonymous substitution rates (dN/dS) to McDonald–Kreitman-style approaches [2] that weigh mutations fixed along a virus’s successful trunk lineage against those lost on unsuccessful side branches [3]. Such approaches have revealed rapid, continued adaptation in the SARS-CoV-2 spike S1 subunit [4, 5].

A complementary class of methods estimates fitness directly from the dynamics of variant frequencies rather than from the composition of mutations [6]. Multinomial logistic regression (MLR) models the frequencies of co-circulating variants through time and infers a relative growth rate, or fitness, for each [7, 8]. Because it expresses fitness as a difference in growth rate between variants, this measure maps directly onto the population-genetic notion of selective advantage. These growth-rate differences correspond to differences in the time-varying effective reproduction number between co-circulating variants [9]. Aggregating these per-variant fitnesses into the rate of change of mean population fitness yields the population’s fitness flux [10], a direct, frequency-based alternative to dN/dS for quantifying the tempo of adaptation.

Here we use this frequency-based view of fitness to trace how SARS-CoV-2 has adapted from the early pandemic in 2020 through 2025, spanning the transition from initial host adaptation to sustained evolution for antigenic novelty. We place the rate of SARS-CoV-2 fitness change in context by comparing it against seasonal influenza A/H3N2 which exhibits canonically rapid and continuous adaptation [11]. Finally, we relate the inferred changes in fitness to molecular predictors, most directly the accumulation of spike mutations, to identify the substitutions that drive fitness gain.

## Results and discussion

### Frequency dynamics

We follow population genetics first principles to compute the frequency through time of a haploid allele under deterministic selection in a fixed-generation Wright–Fisher population. If an allele is at frequency *x*, then after a single generation with selective advantage *s* the expected allele frequency will be

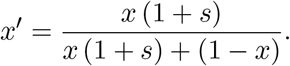

Compounded over *t* generations, the expectation from initial frequency *p* follows

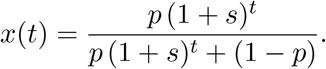

Generalizing this two-allele model to *n* co-circulating variants, each with initial frequency *p*_*i*_ and selective advantage *s*_*i*_, variant *i*’s frequency is its relative abundance normalized by the sum across all variants,

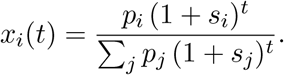

Moving from many discrete generations to continuous time, (1 + *s*_*i*_)^*t*^ *≈*exp(*t* log(1 + *s*_*i*_)), so writing the growth rate *f*_*i*_ = log(1 + *s*_*i*_) gives the probability that a virus sampled at time *t* is labeled as variant *i*

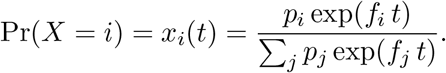

This is the multinomial logistic regression (MLR) model, which has been widely used for modeling SARS-CoV-2 variant frequencies [7, 8]. The denominator normalizes the exponential growth/decay of individual variants so that overall frequency sums to 1. The model has 2*n* parameters, with each variant *i* having an initial frequency *p*_*i*_ and a fixed growth rate *f*_*i*_. Because growth rates are necessarily relative, we fix an arbitrary “pivot” variant as a reference with growth rate *f* = 0. MLR growth rates are directly estimated in terms of calendar time with per-day or per-year values of *f*. To express fitness in per-generation units we multiply each per-day rate by the generation time *τ* measured in days, giving 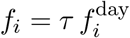, the change in log frequency accrued over a single generation. We assume *τ* of 5.0 days for pre-Omicron SARS-CoV-2, 3.2 days for post-Omicron SARS-CoV-2 and 3.2 for influenza H3N2 (see Methods). Throughout, we refer to this per-generation growth rate *f*_*i*_ = log(1 + *s*_*i*_) as the fitness of variant *i*, or equivalently exp(*f*_*i*_) = 1 + *s*_*i*_ is its pergeneration multiplicative fitness, the factor by which the variant’s abundance grows each generation relative to the pivot. Because *f*_*i*_ is defined on a log scale, mean fitness, fitness variance, fitness flux and changes in fitness between lineages are all likewise computed on this log scale.

We estimate frequencies and fitnesses of SARS-CoV-2 clades in 1-year sliding windows between Jan 2020 and Dec 2025 (Figure 1). In each window we collect clade sequence counts for viruses sampled from the USA and estimate per-variant frequencies and fitnesses. We use only the USA, the one country with sufficient temporal sequencing coverage over this period. We collapse rare clades into a single “other” clade for MLR analysis to prevent noisy estimates from low sequence counts (see Methods). The match between the empirical frequencies (dotted trajectories) and MLR frequencies (solid trajectories) indicates the model fits well despite having few parameters.

**Figure 1.**
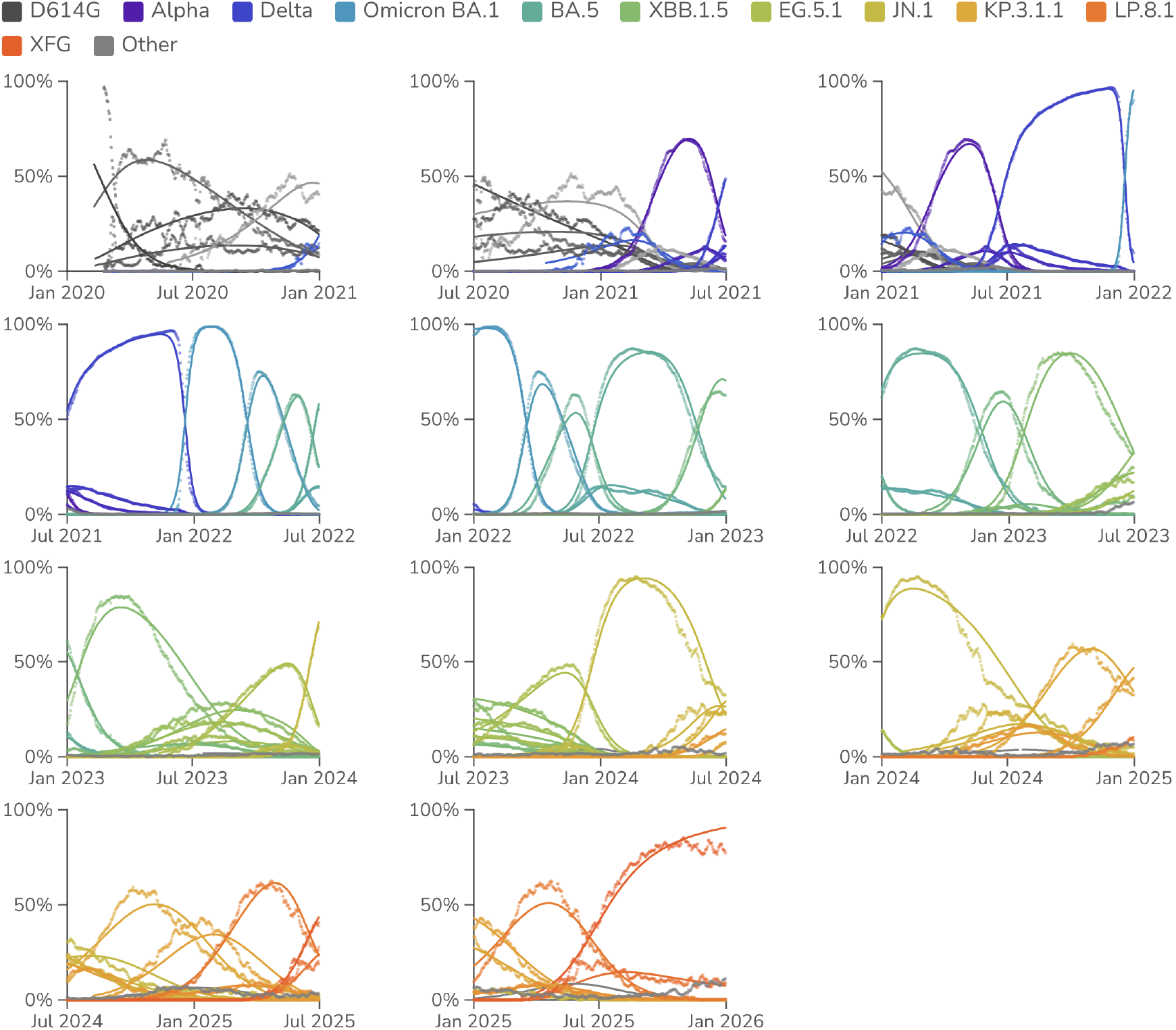
Relative frequencies of SARS-CoV-2 clades through time. Points represent empirical frequencies of SARS-CoV-2 Nextstrain clades, while solid lines represent modeled frequencies from multinomial logistic regression (MLR). All data is taken from the USA. The MLR analysis assumes that the fitness of each clade is constant through time. MLR frequency lines are drawn where there is sufficient sequence data to estimate empirical frequencies.

For comparison purposes, we take a similar approach to estimating frequencies and fitnesses of seasonal influenza H3N2 (Figure 2). Here we use 2-year sliding windows to account for slower frequency dynamics in seasonal influenza and still only use data from the USA. The model fits are worse for H3N2 compared to SARS-CoV-2. This is especially apparent at junctions between influenza seasons where stochastic seeding of a new season may result in a discontinuity of clade frequency compared to MLR expectation. However, H3N2 fits remain sufficient to estimate the magnitude of fitness effects.

**Figure 2.**
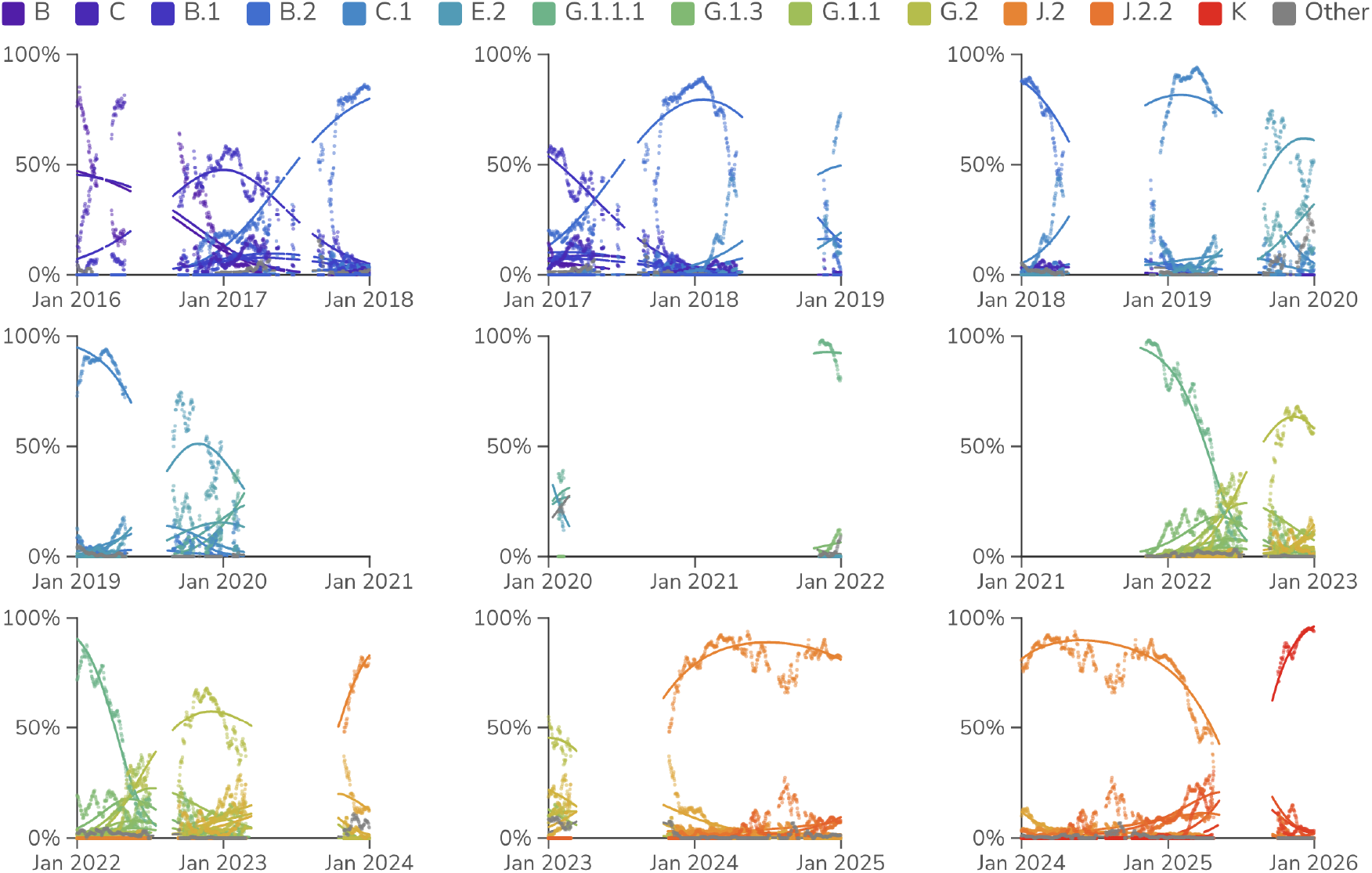
Relative frequencies of H3N2 clades through time. Points represent empirical frequencies of H3N2 Nextstrain clades, while solid lines represent modeled frequencies from multinomial logistic regression (MLR). All data is taken from the USA. The MLR analysis assumes that the fitness of each clade is constant through time. MLR frequency lines are drawn where there is sufficient sequence data to estimate empirical frequencies.

We “scaffold” MLR fitness estimates across windows to arrive at a single fitness estimate per variant. Here, each window only measures fitness differences, with its own arbitrary zero. We solve for the single set of clade fitnesses and per-window offsets that best fit every window at once, weighting by abundance (see Methods).

#### Fitness flux

With variant frequency *x*_*i*_(*t*) and constant variant fitness *f*_*i*_, we describe the mean population fitness as a standard weighted sum 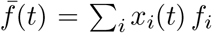. The fitness flux [10] of the population is then the rate of change of population fitness at a given time 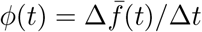.

Integrating this rate gives the cumulative fitness flux

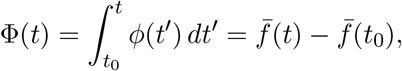

the total adaptive change accumulated along the population’s trajectory. Because variant fitnesses are estimated only relative to a pivot, an individual variant’s scaffolded fitness is meaningful as a difference from a baseline rather than as an absolute value. Chaining these locally-measured advantages across overlapping windows places variant *i* at a cumulative fitness flux Φ_*i*_ = *f*_*i*_ − *f*_0_ relative to the founding variant, and the population sits at the frequency-weighted average Φ(*t*) = ∑_*i*_ *x*_*i*_(*t*) Φ_*i*_.

We find that SARS-CoV-2 initially accumulated fitness flux rapidly with mean fitness doubling every 6 months between Jan 2021 and Jun 2022, but then slowing to doubling every 2.4 years from July 2022 to Dec 2025 (Figure 3). After initial spread of D614G [12] in 2020, we observe a lull, followed by rapid growth in fitness in 2021 and 2022 with initial VOCs, Omicron and initial Omicron sub-lineages [13], and then a slower, more steady pace since 2024. There is a mix of large jumps in fitness (familiar examples like Delta and BA.1, but also more recently with JN.1) and smaller, more gradual step change.

**Figure 3.**
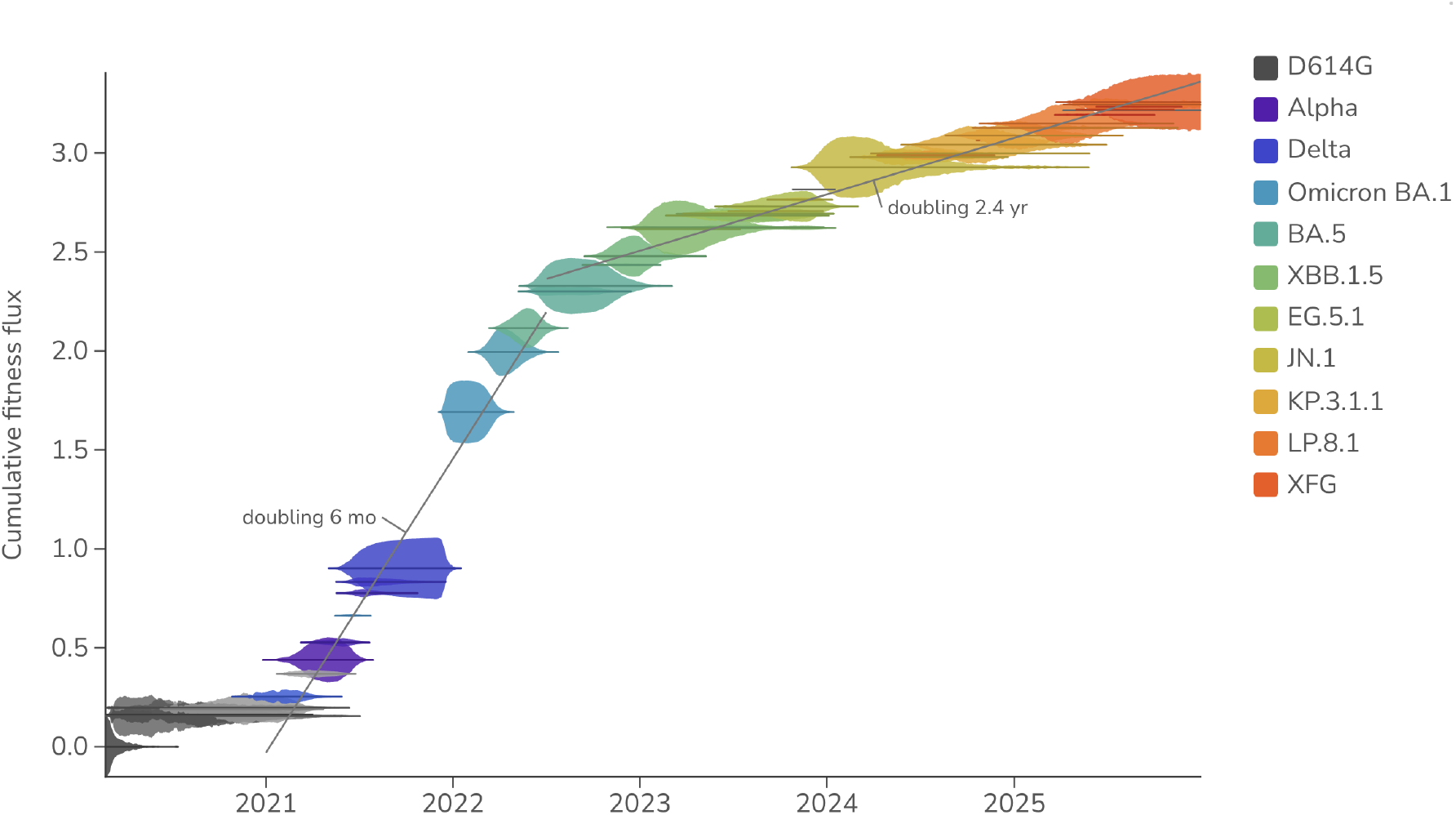
Cumulative SARS-CoV-2 fitness flux. Empirical frequencies of SARS-CoV-2 clades are represented by vertical thickness and placement on the y-axis represents cumulative fitness flux estimated from multinomial logistic regression (MLR). Gray lines are least-squares fits to the mean fitness over each labeled period, annotated with the implied doubling time. All data is taken from the USA. The MLR analysis assumes that the fitness of each clade is constant through time.

Seasonal influenza H3N2 shows a fundamentally similar pattern of emergence of new clades and their replacement of existing diversity. However, H3N2 dynamics play out on a slower timescale (Figure 4). Rather than SARS-CoV-2’s months-scale doubling, H3N2 adapts much more slowly, doubling roughly every 10.0 years over the course of 2016 to 2025. Greater coexistence of multiple co-circulating clades is also apparent relative to SARSCoV-2.

**Figure 4.**
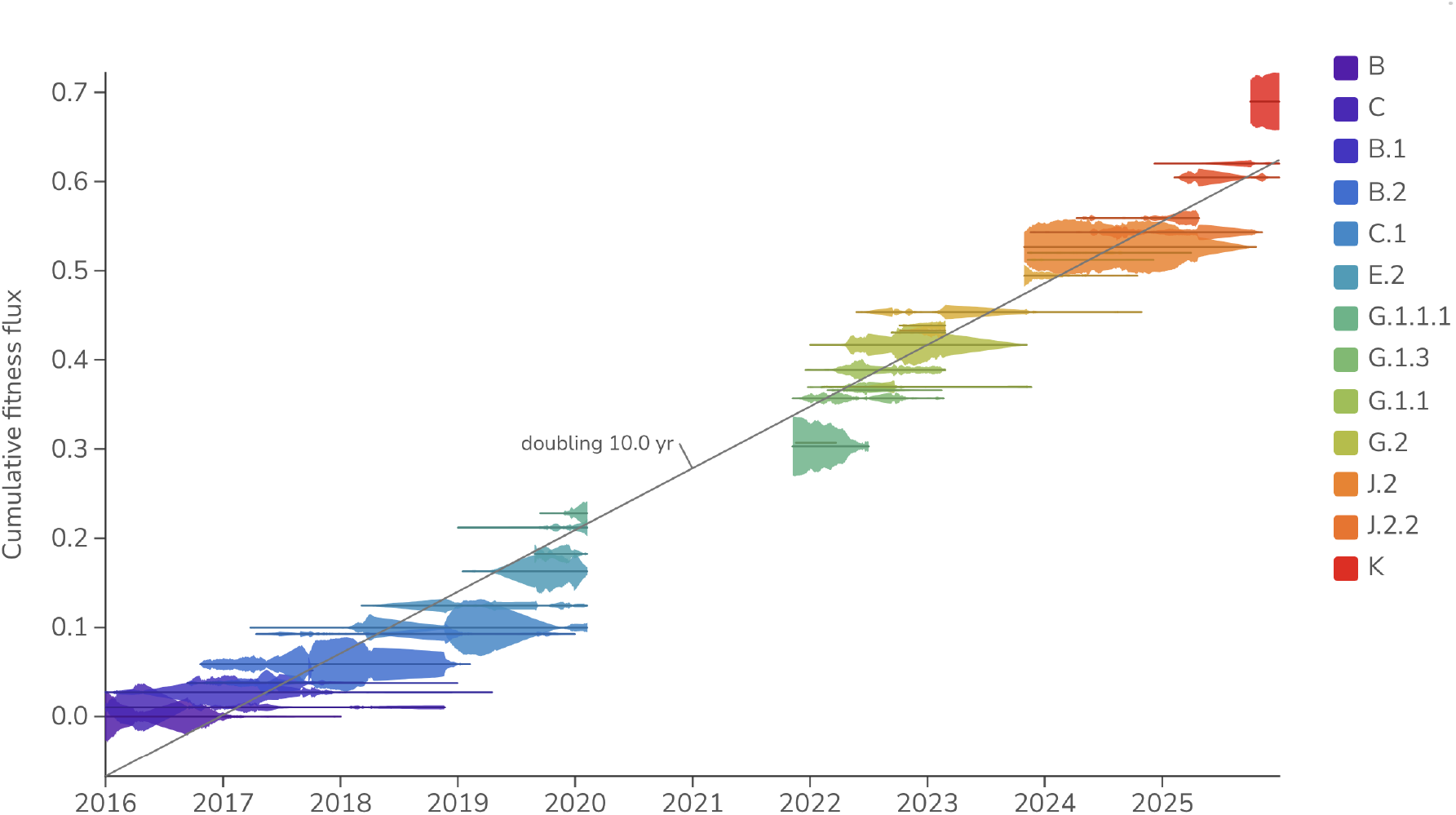
Cumulative H3N2 fitness flux. Empirical frequencies of H3N2 clades are represented by vertical thickness and placement on the y-axis represents cumulative fitness flux estimated from multinomial logistic regression (MLR). Gray lines are least-squares fits to the mean fitness over each labeled period, annotated with the implied doubling time. All data is taken from the USA. The MLR analysis assumes that the fitness of each clade is constant through time.

SARS-CoV-2 clade frequencies and fitnesses can be viewed as a phase portrait, plotting each clade’s empirical frequency against its fitness relative to the daily population average (Figure 5). A clade emerges at low frequency and high relative fitness, sweeps up in frequency as its relative fitness declines toward the population average, peaks near a relative fitness of zero, and then falls back to low frequency as it is outcompeted. Clades that start out with a greater advantage over the population average tend to sweep to higher maximum frequency than clades that start with less of an advantage.

**Figure 5.**
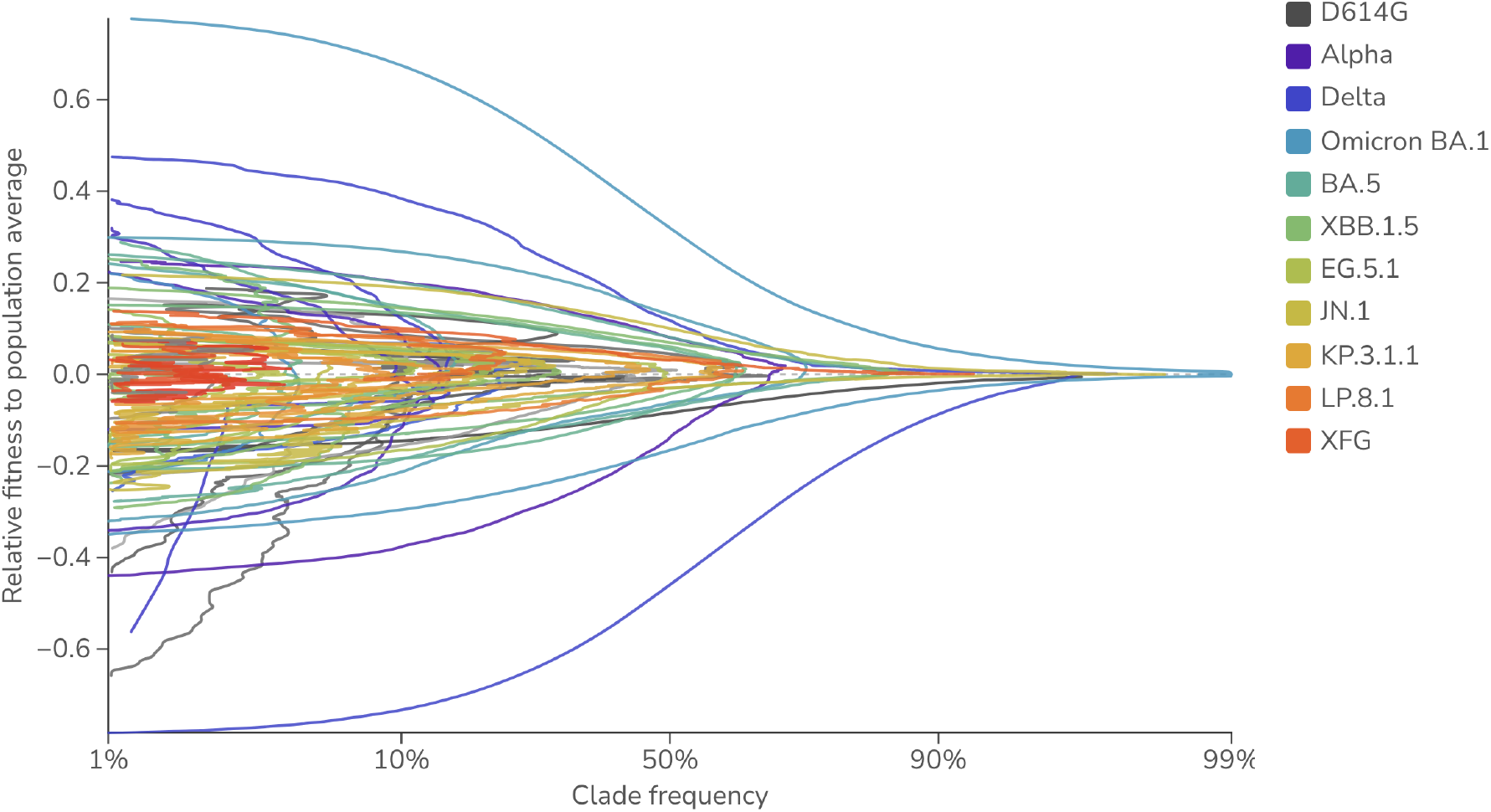
Frequency vs fitness phase portrait for SARS-CoV-2 clades. Each line traces a SARS-CoV-2 clade’s trajectory over time through empirical frequency (x-axis, logit scale) and fitness relative to the daily population average (y-axis), estimated from multinomial logistic regression (MLR). All data is taken from the USA. The MLR analysis assumes that the fitness of each clade is constant through time.

#### Fisher’s fundamental theorem

Multistrain models that allow for antigenic evolution produce traveling waves in antigenic space [14]. More broadly, many mutations of small fitness effect create traveling fitness waves where the rate of advance toward higher fitness is proportional to the variance in fitness [15]. This is a consequence of Fisher’s fundamental theorem of natural selection

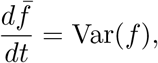

where “the rate of increase in fitness of any organism at any time is equal to its genetic variance in fitness at that time” [16].

We can investigate this relationship directly in SARS-CoV-2 (Figure 6), where we find that timepoints with larger variance in fitness Var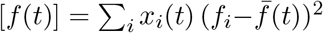 correlate well with timepoints with larger change in mean population fitness 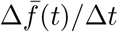. In fact we find that the relationship is near the 1:1 expectation from Fisher’s theorem (slope = 1.10, Pearson *r* = 0.94). Looking in detail at rate of fitness flux through time, we find a yearly average fitness flux of 18.0 ×10^−3^ per-gen in 2021 followed by a reduction to 1.7 − 1.8 × 10^−3^ per-gen in 2024 and 2025. This shows that the rate of adaptation of SARS-CoV-2 has been slowing as low hanging fruit of host adaptation is exhausted, leaving only red-queen antigenic evolution to drive adaptation.

**Figure 6.**
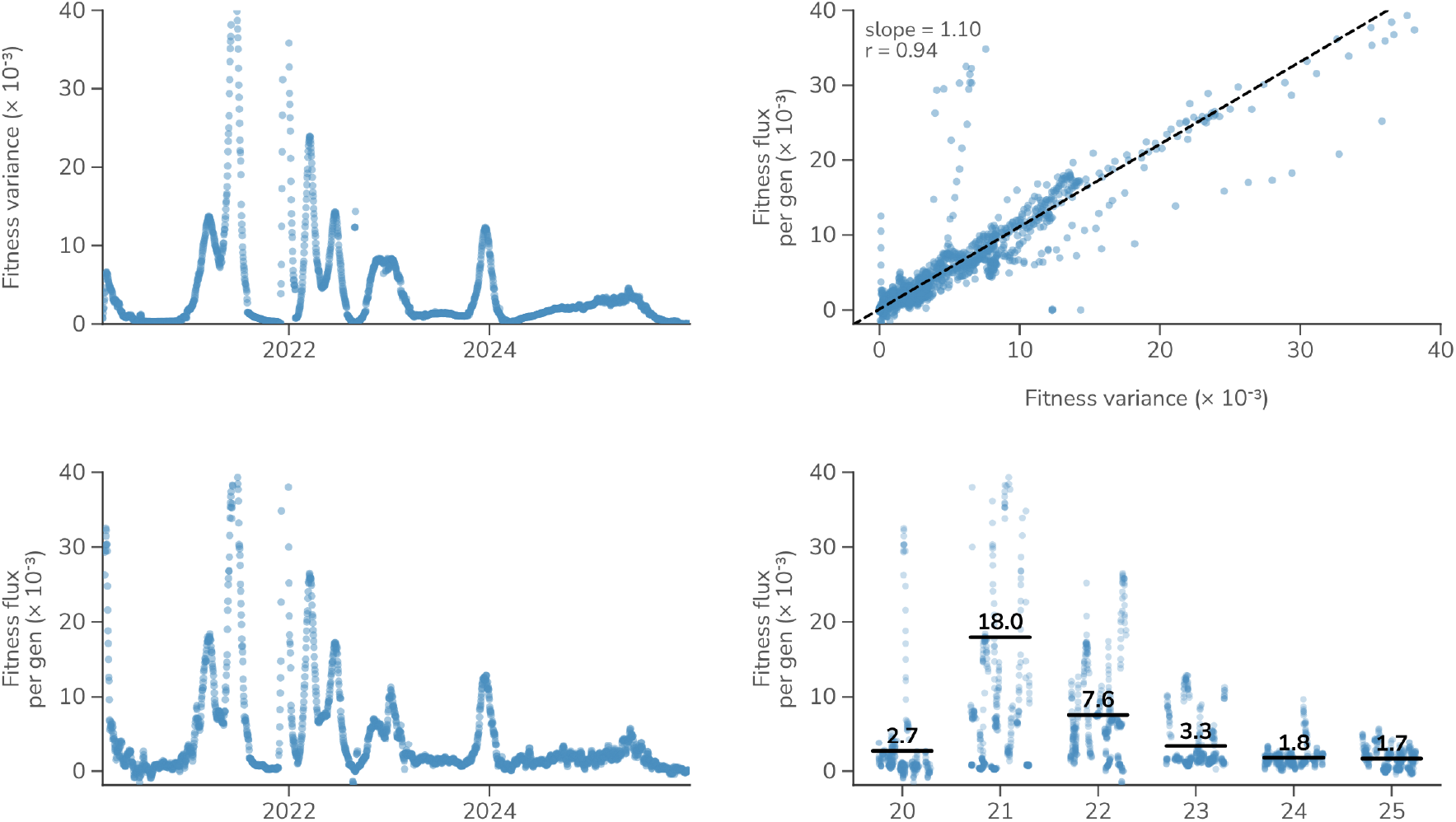
Fitness variance and fitness flux in SARS-CoV-2. Fitness variance is compared to fitness flux, where each dot represents a daily timepoint.

Compared to SARS-CoV-2, influenza H3N2 shows generally lower rates of fitness flux, averaging 0.5 × 10^−3^ per-gen from 2016 to 2025 (Figure 7). This is roughly 3 times lower than recent years of SARS-CoV-2 fitness flux. However, it remains possible that SARSCoV-2 slows further in the coming years.

**Figure 7.**
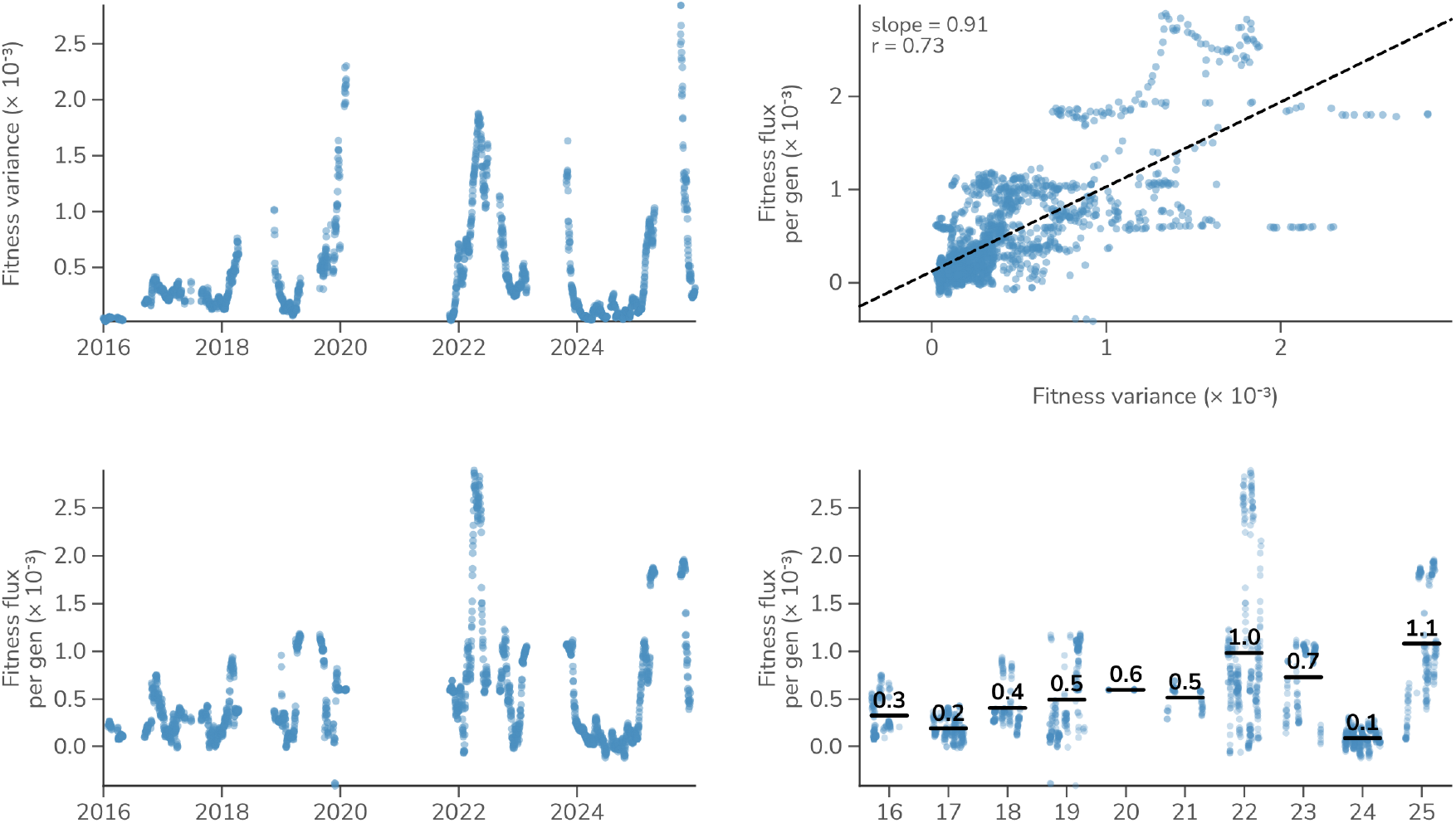
Fitness variance and fitness flux in H3N2. Fitness variance is compared to fitness flux, where each dot represents a daily timepoint.

For both SARS-CoV-2 and influenza H3N2, the connection between fitness flux and fitness variance is clear. This suggests from first principles that interventions that decrease variance in fitness across the virus population would be expected to slow adaptation, while interventions that increase variance would be expected to speed adaptation.

#### Mutational fitness effects

A simple correlation of cumulative mutations against cumulative fitness flux will fail due to phylogenetic non-independence, and instead we rely on phylogenetic contrasts of parent and child lineages [17]. Pango lineages [18] provide a convenient granular and hierarchical nomenclature well suited to this.

We define Pango parent-to-child branches by finding mutations that accumulate between hierarchical Pango lineages. Given estimates of per-lineage fitness, we also calculate the change in fitness between each parent and child. As expected, we observe that Pango lineages are granular with parent-to-child changes in spike mutations, non-spike mutations and fitness being modest, with most branches adding only a handful of substitutions and shifting fitness by a small amount (Figure 8).

**Figure 8.**
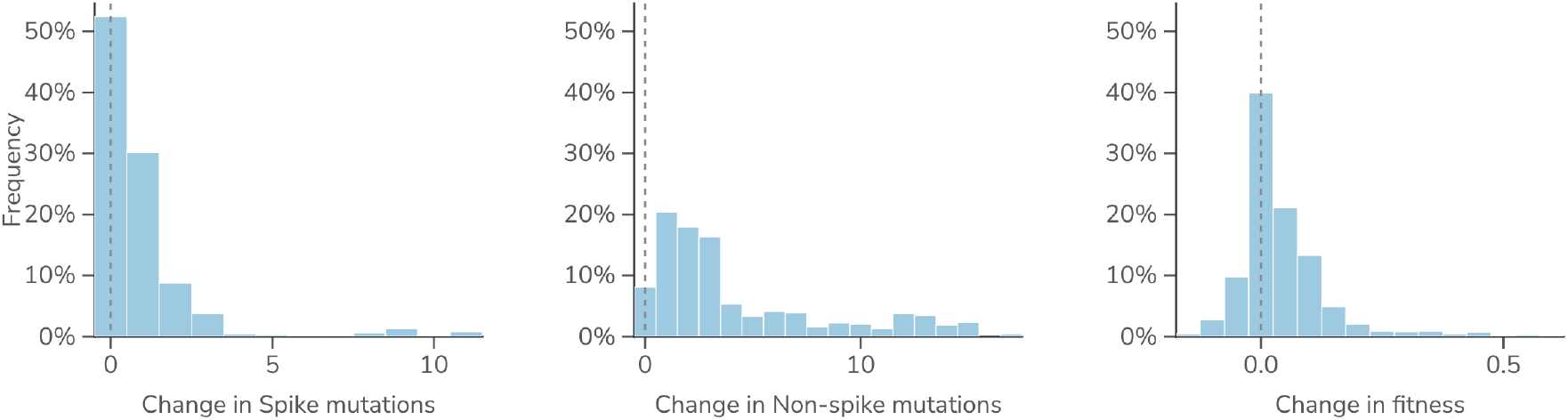
Distributions of mutation and fitness change across SARS-CoV-2 lineage branches. Across all parent-to-child Pango lineage branches, the change in the number of spike substitutions, the change in non-spike substitutions, and the change in fitness. Each bar is the fraction of branches in that bin; rare large founder jumps fall beyond the plotted range.

We compare the change in fitness along each parent-to-child branch against the substitutions it acquired in different regions of the SARS-CoV-2 genome (Figure 9). Spike S1 substitutions and those in the receptor-binding domain (RBD) in particular, carry the strongest positive association with fitness gain, while ORF1ab and accessory gene substitutions show lower regression slopes and correlation coefficients. Splitting branches into early and late periods shows that the fitness value of a given substitution is largest early and decays over time.

**Figure 9.**
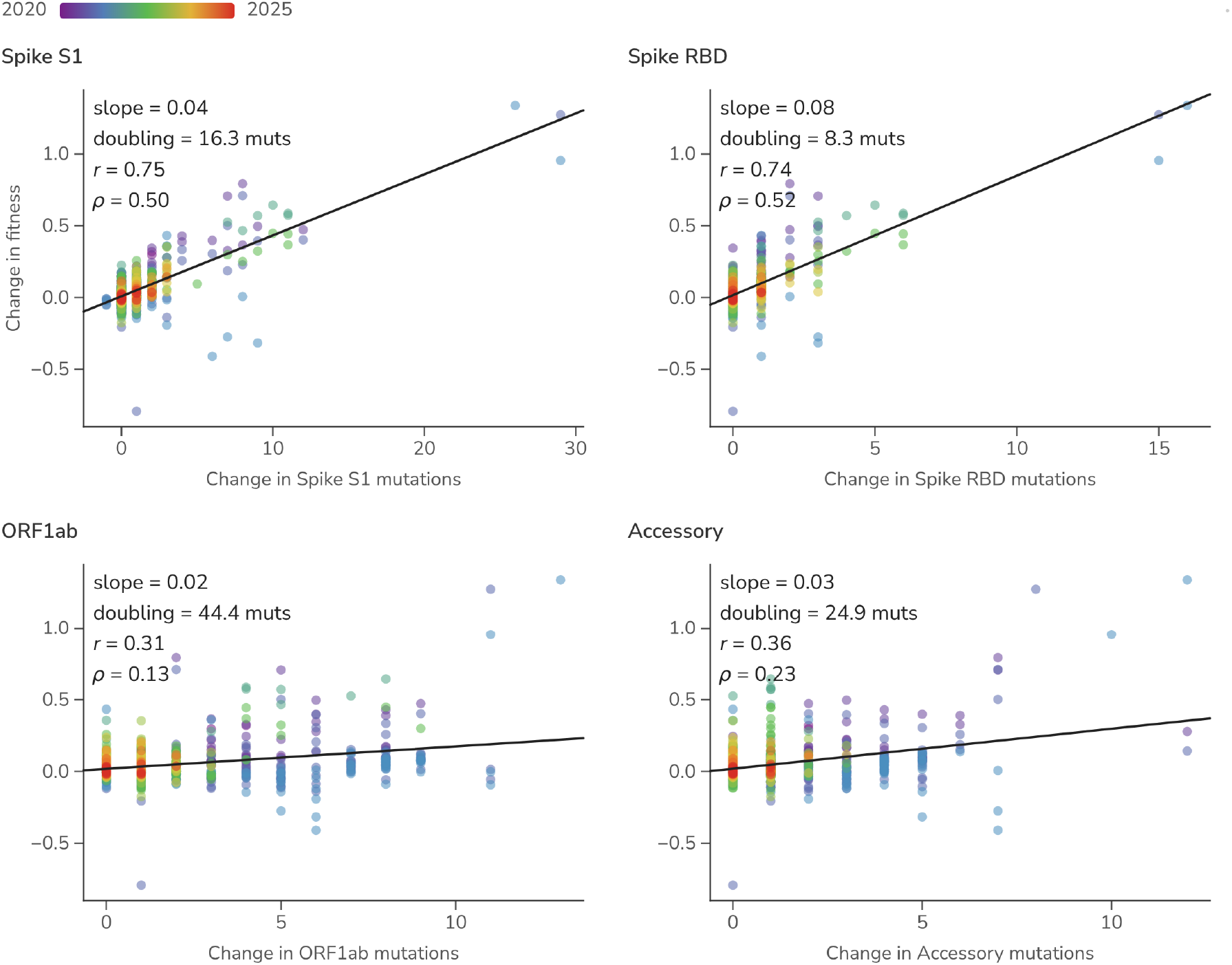
Lineage-specific amino acid change versus lineage-specific fitness change across regions of the SARS-CoV-2 genome. Each point is one parent-to-child Pango lineage branch in one season: the change in the number of substitutions in a genome region (x) against the change in fitness (y), colored by time from blue (2020) to red (2025), with a least-squares fit per panel.

We can track this relationship through time by fitting the regression separately within each season (Figure 10). The per-substitution effect on fitness of spike and RBD changes is largest early and erodes toward zero as the readily accessible routes to host adaptation are exhausted, while other regions stay near zero throughout. However, across accessory proteins and in ORF1ab there is moderate marginal correlation from 2020 to 2022 between substitutions and fitness change; the multiple regression below shows this to be a confound of co-occurrence with spike rather than an independent effect. Although per-substitution effect (ie regression slope) of RBD decays from 2020, the predictive ability of spike S1 and RBD substitutions as measured by Pearson and Spearman correlations stays high through the period with average correlation coefficients of *r* = 0.72 and *ρ* = 0.54 for spike S1 and *r* = 0.70 and *ρ* = 0.53 for spike RBD.

**Figure 10.**
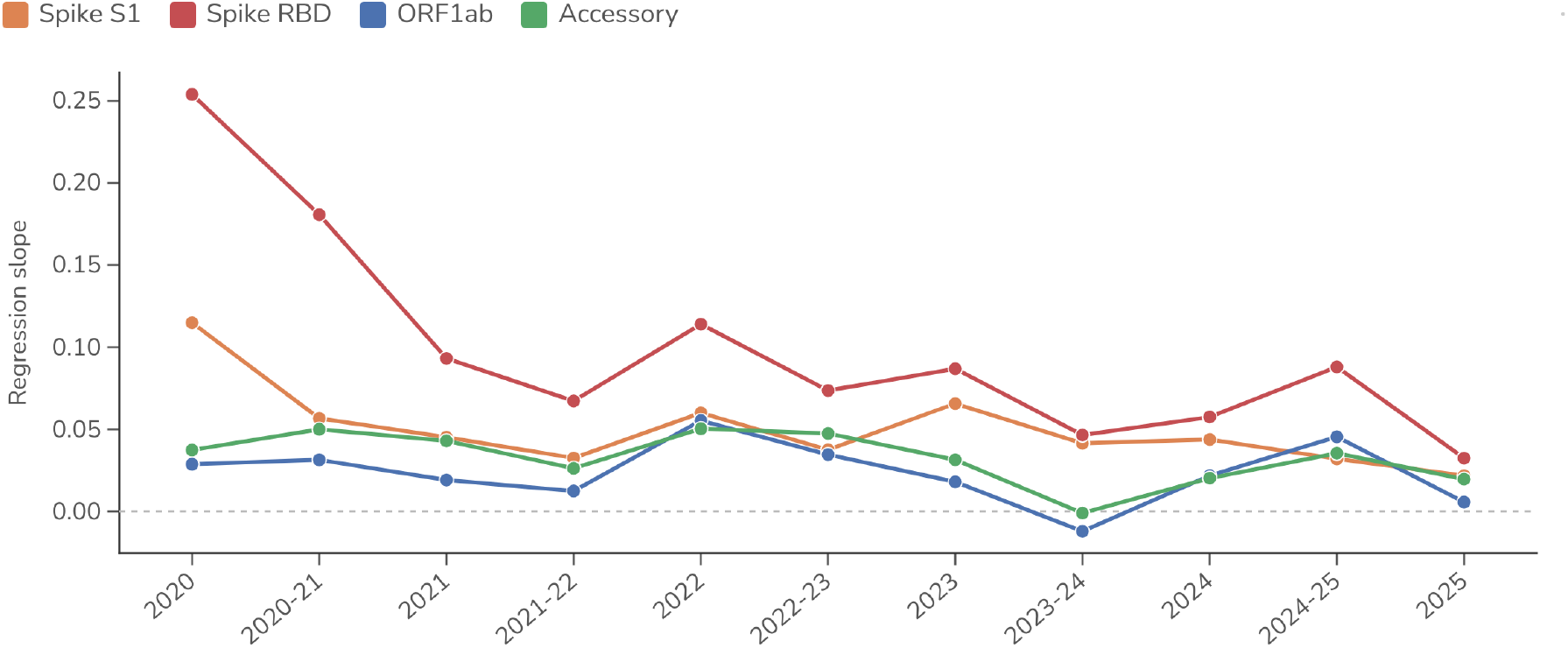
Strength of the mutation–fitness relationship through time. For each season the parent-to-child branches are summarized by regression slope relating change in regional substitutions to change in fitness, with one line per genome region.

The marginal relationships in Figure 9 cannot on their own establish which substitutions drive fitness. A more evolved lineage accumulates more substitutions across the whole genome, so a region can correlate with fitness merely by co-varying with a genuinely causal region. The moderate marginal association of ORF1ab substitutions is a case in point.

To isolate each region’s independent contribution we fit a multiple linear regression of the change in substitution count across four non-overlapping genome regions to per-branch change in fitness (Figure 11). Once spike is controlled for, most of the positive signal sits in the RBD (*β* = 0.064 per substitution, *p* < 10^−20^), with a smaller contribution from the remainder of S1 (*β* = 0.015, *p* = 1.5 ×10^−4^), while the ORF1ab and accessory-protein coefficients collapse to near zero and are no longer distinguishable from no effect (*β* = 0.002, *p* = 0.32 and *β* = 0.003, *p* = 0.41, respectively). The apparent marginal association of ORF1ab is therefore a confound of its co-occurrence with spike change rather than evidence that ORF1ab substitutions themselves raise fitness. The four-region model explains most of the variance in per-pair fitness change (*R*^2^ = 0.73), with predicted and observed changes falling along the 1:1 line.

**Figure 11.**
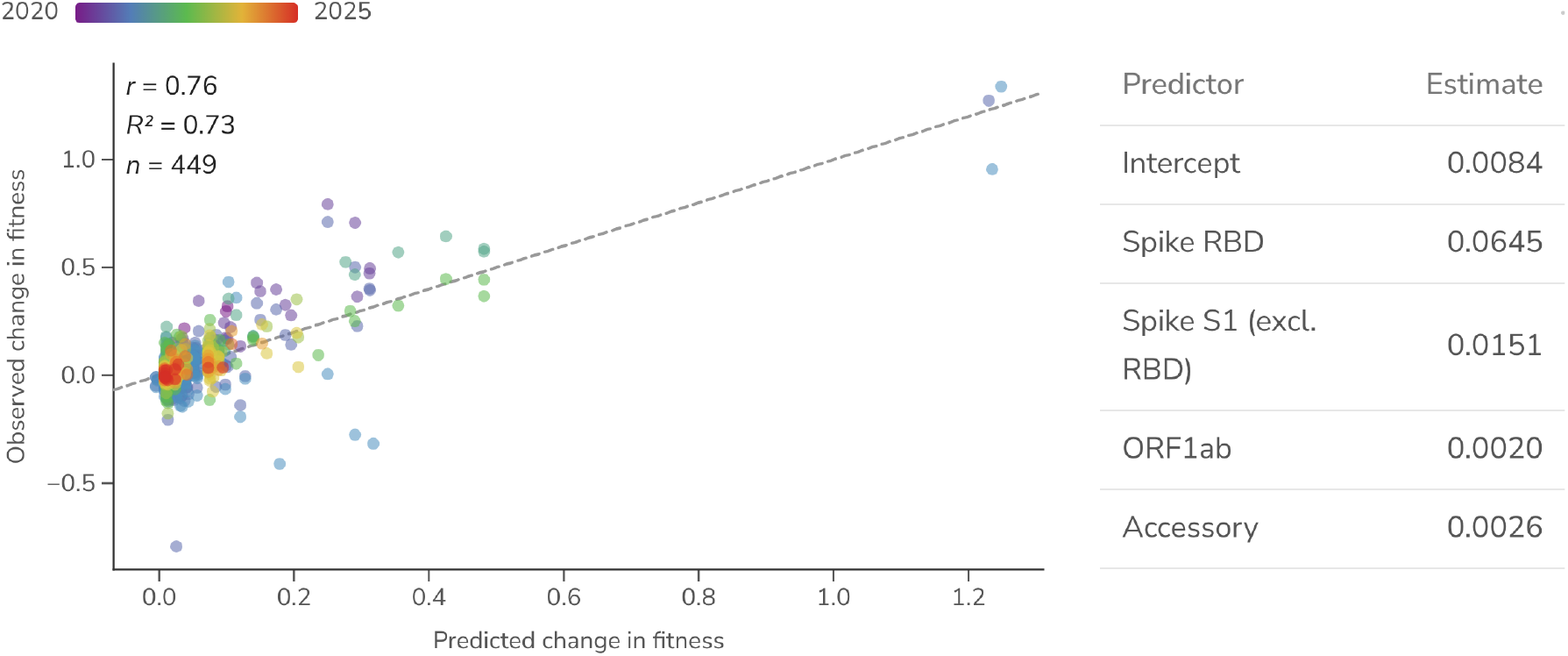
Multiple regression of fitness change on non-overlapping genome regions. An ordinary-least-squares fit of the change in fitness on the change in substitution count in four non-overlapping regions of the SARS-CoV-2 genome (spike RBD, spike S1 outside the RBD, ORF1ab, accessory), fit over unique parent-to-child pairs (collapsing observations of a pair that recur across windows). The table gives each region’s partial estimate. The scatter plots each parent-to-child Pango lineage branch’s model-predicted change in fitness (x) against its observed change in fitness (y), colored by time from blue (2020) to red (2025) and with dashed 1:1 calibration line.

#### Predicting fitness effects

The core idea of comparing change in mutation count to change in fitness expresses a similar logic to McDonald-Kreitman tests [2] where the key comparison is relative success of lineages bearing different mutation patterns. This predictor-vs-growth-rate formulation [4] should be robust to many confounders that affect other measures of adaptation.

We compare our simple, non-parameterized predictor of relative spike S1 substitution count against two recent deep-learning proposals for scoring evolutionarily successful substitutions (Figure 12). These are EvEscape [19] and protein-language-model semanticity [20], the latter reimplemented with pretrained and fine-tuned ESM-2 embeddings [21] (see Methods). ESM is fine-tuned to SARS-CoV-2 spike sequences from 2020 through 2022. Both perform similarly to a plain count of spike S1 substitutions in disambiguating the fitness of co-circulating SARS-CoV-2 lineages.

**Figure 12.**
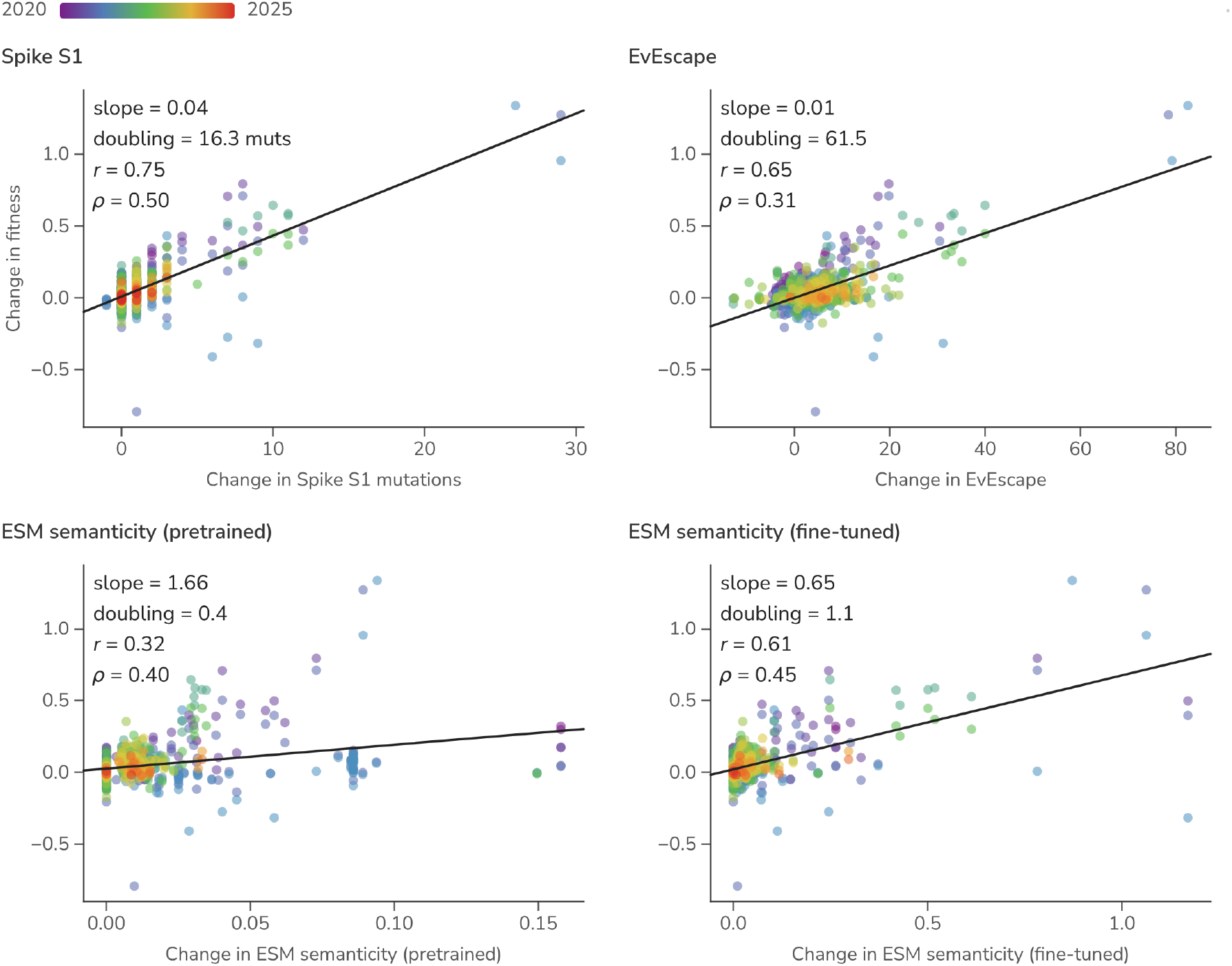
Lineage-specific predictors versus lineage-specific fitness change. Each point is one parent-to-child Pango lineage branch in one season: change in predictor value (x) against the change in fitness (y), colored by time from blue (2020) to red (2025), with a least-squares fit per panel.

## Conclusions

Most measures of viral adaptation are indirect, diagnosing the presence of selection based on mutation patterns. Here we instead read adaptation directly off the dynamics of variant frequencies, aggregating per-variant growth rates into the population’s fitness flux. This turns the tempo of adaptation into a single quantity that can be followed through time, placed on a common per-generation scale across pathogens and connected to first principles. On this scale SARS-CoV-2 adapts rapidly, doubling in fitness roughly every 6 months during initial variant emergence in 2021 and early 2022, but decelerating towards a baseline flux in 2024, while seasonal H3N2 sustains a slower, steadier flux. Importantly, these numbers carry an absolute scale and connect back to epidemiological impact [9].

The generality of the approach rests on a single requirement: a way to bin genetic diversity into discrete, comparable variants. For SARS-CoV-2 and influenza this comes off the shelf, with Nextstrain clades supporting the frequency and flux analysis and finer, hierarchically nested Pango lineages supporting the phylogenetic-contrast analysis of mutational effects. Pathogens without an established nomenclature could be analyzed via automated methods that partition a tree into lineages [22, 23].

Beyond describing historical adaptation, the per-branch contrast of mutation against fitness change yields a simple, interpretable account of which substitutions matter. For SARSCoV-2 the signal concentrates in spike (and particularly spike RBD) and a plain count of spike S1 substitutions disambiguates the relative fitness of co-circulating lineages about as well as recently proposed deep-learning escape and protein-language-model scores [19, 20]. This makes the mutation-to-fitness deltas a strong and transparent baseline for forecasting variant success, where a predictor that does not improve on counting spike substitutions has not yet justified its added complexity.

## Methods

### Sequence data

For SARS-CoV-2, we use curated “open” data from Nextstrain [24] that draws from NCBI GenBank. For influenza H3N2, we use data from GISAID [25]. In each case, the raw sequences are processed with Nextclade [26] to assign Nextstrain clade and Pango lineage to [18] SARS-CoV-2 sequences and to assign subclade [27] to influenza H3N2 sequences. We filter out sequences with Nextclade overall QC status of “bad”. We additionally filter to sequences collected from the USA. This leaves 3,588,802 total sequences for SARS-CoV-2 sampled between 2020 and 2025 and 44,456 total sequences for H3N2 sampled between 2016 and 2025.

### Multinomial logistic regression

We conducted multinomial logistic regression (MLR) using the evofr package (github.com/blab/evofr) on 1-year sliding windows for SARS-CoV-2 (11 windows total) and 2-year sliding windows for H3N2 (9 windows total). For each window we treat each clade as a distinct variant, collapsing rare clades together into a single “other” category before fitting. For both SARSCoV-2 and H3N2, a clade is modeled separately only if it reaches at least 50 sequences and a mean frequency of at least 0.1% within the window, while clades below either threshold are merged into “other”. This leaves between 7 and 18 clades per window (median 15) for SARS-CoV-2 and between 5 and 13 (median 9) for influenza H3N2.

We use generation time *τ* of 5.0 days for pre-Omicron SARS-CoV-2 following [28–30], generation time of 3.2 days for post-Omicron SARS-CoV-2 following [31,32] and generation time of 3.2 days for seasonal influenza H3N2 following [33–35]. At first order, the fitness 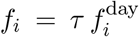 is the log ratio of reproduction numbers between variant *i* and the pivot, log(*R*_*i*_*/R*_pivot_). Only the mean generation interval *τ* enters at this order. The shape of the generation-interval distribution enters at second order, informed by the interval’s variance and absolute per-day growth rates rather than 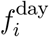. Because the frequency data identify only this relative rate, the second order correction cannot be estimated from frequency dynamics alone. The per-generation multiplicative fitness exp(*f*_*i*_) = 1 + *s*_*i*_ therefore equals the reproduction-number ratio *R*_*i*_*/R*_pivot_ under a fixed (point) generation interval and is an upper bound under a realistically dispersed interval [36], for which the fuller frequency-to-reproduction-number conversion is given in Figgins and Bedford [9].

We conduct a parallel MLR analysis of SARS-CoV-2 Pango lineages. Because lineages are hierarchically nested, rather than collapsing rare lineages into a shared “other” we roll each lineage with fewer than 500 sequences up into its parent lineage, repeating until every retained lineage clears this count. A lineage is additionally retained only if at least 200 sequences are assigned to that lineage itself rather than to a descendant sub-lineage, otherwise it is folded into “other”. This leaves between 20 and 165 lineages per window (median 79) for SARS-CoV-2. Rationale for specific collapse cutoffs is available at github.com/blab/fitness-flux/tree/main/inclusion-thresholds.

### Scaffolding across timepoints

Within each sliding window the MLR model estimates each variant’s fitness only relative to that window’s pivot, so every window carries its own arbitrary additive zero and the per-window estimates *f*_*i,w*_ are not directly comparable. We recover a single fitness per variant by treating scaffolding as a weighted two-way additive model: each estimate is a variant effect minus a window effect, *f*_*i,w*_*≈ f*_*i*_ −*c*_*w*_, where *f*_*i*_ is variant *i*’s global fitness and *c*_*w*_ is window *w*’s offset. We choose the *f*_*i*_ and *c*_*w*_ that jointly minimize the abundance-weighted squared error across every window,

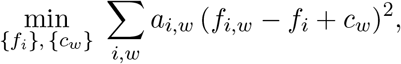

weighting each estimate by *a*_*i,w*_, the area under variant *i*’s modeled-frequency curve in window *w*, so a window in which a variant is rare with poorly constrained MLR estimate will contribute negligibly. The optimum is a pair of interleaved abundance-weighted means, each variant’s fitness being the weighted mean of its offset-corrected estimates over its windows

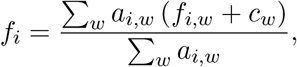

and each window’s offset the weighted-mean gap between the global scale and that window’s estimates

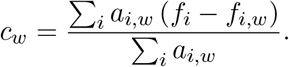

We solve by alternating the two to convergence. The overlap of variants between windows ties them into one connected scale, leaving a single global constant free, which we fix by shifting all values so that the founding variant, our least-fit baseline, sits at zero, leaving each variant’s scaffolded value as its cumulative fitness flux Φ_*i*_ = *f*_*i*_ − *f*_0_.

### Lineage mutation counts and branch contrasts

To relate change in fitness to change in genotype, we count amino-acid substitutions per Pango lineage and compare each lineage against its parent. Per-lineage substitution counts are read from the Nextclade SARS-CoV-2 reference tree at nextstrain.org/nextclade/nextstrain/sars-cov-2/wuhan-hu-1/orfs, in which each tip corresponds to a Pango lineage. For each lineage we count substitutions relative to the Wuhan-Hu-1 reference. We tally these by region: the spike S1 subunit, the receptor-binding domain (RBD, 319–541) within S1, ORF1ab, and the accessory and structural genes (ORF3a, E, M, ORF6, ORF7a, ORF7b, ORF8, N).

We then form parent-to-child branches between hierarchically nested Pango lineages. Within each window a lineage’s parent is its closest retained ancestor, where the retained set is fixed by the collapsing described above, so lineages that were rolled up into a parent or folded into “other” do not themselves appear as branch endpoints. For every branch whose parent and child both carry an MLR fitness estimate in that window, we record the change in substitution count in each genome region and the change in fitness, taken as the difference in their per-window MLR fitness; because both endpoints are estimated against the same window’s pivot, this contrast is well defined without scaffolding. A branch is recorded once per window in which it appears, so a lineage pair that co-circulates across several windows contributes several observations. These per-branch deltas in mutation count and fitness are the unit of the mutational-fitness analyses. For the multiple regression we collapse these repeated observations to a single contrast per parent-to-child pair, averaging the per-window fitness change, so that coefficients and *p*-values are not inflated by this pseudoreplication.

### Mutational fitness predictors

We compare the per-branch substitution counts above against two externally proposed predictors of mutational fitness, each reduced to a per-lineage value and contrasted across the same parent-to-child branches. EvEscape [19] combines a variational-autoencoder fitness model with residue accessibility and biochemical dissimilarity to score spike mutations. We use the precomputed all-strain EvEscape scores released at evescape.org/data and take each Pango lineage’s score as the mean EvEscape across the sequences assigned to that lineage.

Semanticity follows the semantic-change measure of Hie et al. [20], reimplemented with ESM-2 [21]. Each Pango lineage’s spike amino-acid sequence is embedded with the 650M-parameter ESM-2 model (`esm2_t33_650M_UR50D`), taking the CLS-token representation of the final (33rd) layer as a 1280-dimensional sequence embedding. We embed lineages with both the released pretrained weights and weights fine-tuned under a masked-language-model objective (15% of residues masked) for one epoch (AdamW, learning rate 5 × 10^−5^) on roughly 16,000 SARS-CoV-2 spike sequences collected from 2020 through 2022. Finetuning and embedding code is available at github.com/blab/embedded-pathways. The semanticity of a branch is the Euclidean distance between its child and parent lineage embeddings.

### Reproducibility

The workflow to reproduce data, analyses and manuscript is available at github.com/blab/fitness-flux. SARS-CoV-2 sequence data were sourced from publicly available open data in NCBI GenBank and so this side of the workflow is easily runnable. However, influenza H3N2 sequence data was collected from GISAID and these data are not allowed to be reshared and consequently the influenza side of the workflow is unfortunately not reproducible without separately providing GISAID data.

## Acknowledgments

We thank Jumpei Ito, Michael Lässig, Erick Matsen, Richard Neher, Cécile Viboud and members of the Bedford Lab for helpful feedback. SARS-CoV-2 analyses are based on open data in GenBank. We gratefully acknowledge the researchers and data contributors who collected the specimens, generated and deposited the raw sequence data and metadata into NCBI GenBank. Influenza analyses are based on GISAID data. We gratefully acknowledge all data contributors, i.e., the Authors and their Originating laboratories responsible for obtaining the specimens, and their Submitting laboratories for generating the genetic sequence and metadata and sharing via the GISAID Initiative, on which this research is based. TB was funded as a Howard Hughes Medical Institute Investigator. This research was supported in part by grant NSF PHY-2309135, the Gordon and Betty Moore Foundation grant no. 2919.02, and the Chan Zuckerberg Initiative DAF grant to the Kavli Institute for Theoretical Physics (KITP).

